# Neural responses underlying ITD discrimination as a function of sensory reliability in the barn owl

**DOI:** 10.1101/2025.06.10.658945

**Authors:** Brian J. Fischer, Keanu Shadron, Clifford H. Keller, Avinash D.S. Bala, Fanny Cazettes, Roland Ferger, José L. Peña

## Abstract

Discrimination of sensory stimuli is fundamentally constrained by the information encoded in neuronal responses. In the barn owl, interaural time difference (ITD) serves as a primary cue for azimuthal sound localization and is represented topographically in the midbrain auditory space map in the external nucleus of the inferior colliculus (ICx). While prior studies have demonstrated a correspondence between spatial tuning and behavioral acuity, it remains unclear how changes in sensory reliability influence this relationship. Here, we examined how behavioral and neuronal ITD discrimination thresholds vary with binaural correlation (BC), which manipulates ITD cue reliability. Using the pupil dilation response (PDR) as a behavioral metric in head-fixed owls, we found that ITD just-noticeable-differences increased exponentially as BC decreased. In contrast, the widths of ICx ITD tuning curves increased more modestly, indicating that tuning resolution alone does not account for behavioral discrimination performance. By computing the Fisher information from ICx neuronal responses, we showed that the average neuronal discriminability predicts behavioral thresholds across BC levels. A habituation-based model incorporating BC-dependent changes in tuning width, firing rate, and response variability successfully accounted for both direction and ITD discrimination. These findings support a model in which perceptual acuity is governed by the combined influence of neuronal tuning and variability and provide a unified framework for understanding how midbrain auditory representations underlie adaptive spatial hearing.

## Introduction

How does the neural representation of sensory information influence the ability to discriminate changes in sensory stimuli? The information present in neuronal responses is impacted by many factors including the selectivity, magnitude, and variability of responses to sensory signals (Seung and Sompolinsky, 1993; Abbott and Dayan, 1999; Pouget et al., 2003; Harper and McAlpine, 2004; Averbeck et al., 2006; Moreno-Bote et al., 2014). Determining how features of neural responses influence sensory discrimination abilities is essential for understanding how sensory processing translates into perception and behavior, a key question in neuroscience. Here we investigate the relationship between neural coding and sensory discrimination in the barn owl’s sound localization system.

Previous work on sound localization in barn owls supports the hypothesis that the resolution of spatial coding in the midbrain auditory space map determines the owl’s ability to discriminate sound source location (Bala et al., 2003, 2007). Behavioral and neurophysiological studies investigating direction discrimination found a correspondence between the spatial resolution of spatially selective auditory neurons in the external nucleus of the inferior colliculus (ICx) and the minimum audible angle (Bala et al., 2003, 2007). While neuronal spatial resolution accounted for the difference in discrimination performance in azimuth and elevation, these studies did not manipulate the reliability of sensory information by adding noise to the stimulus. The reliability of sensory information is known to affect the ability to discriminate stimuli and influences multiple aspects of neuronal responses (Albeck and Konishi, 1995; Saberi et al., 1998; Peña and Konishi, 2004; Cazettes et al., 2016). Thus, how the reliability of a sound localization cue relates to neuronal and perceptual discrimination in barn owls remains an open question.

Barn owls primarily use the interaural time difference (ITD) to determine the azimuthal direction of a sound source (Moiseff, 1989; Poganiatz et al., 2001; Kettler et al., 2017). The reliability of ITD can be manipulated by changing the binaural correlation (BC), which is achieved by adding independent noise sounds to the left and right ears (Saberi et al., 1998). Prior behavioral experiments with owls localizing sounds by turning the head show that localization accuracy decreases with BC (Saberi et al., 1998). Studies in humans show a similar decrease in localization and discrimination performance as BC decreases (Trahiotis et al., 2001; Rakerd and Hartmann, 2010).

Space-specific neurons are known to exhibit changes in ITD tuning such that the spatial resolution, firing rates, and membrane potential variability decrease as BC decreases (Albeck and Konishi, 1995; Saberi et al., 1998; Rich et al., 2015; Cazettes et al., 2016). Theoretical studies have suggested that the change in ITD tuning resolution can explain the decrease in the owl’s localization accuracy as BC decreases (Fischer and Peña, 2011; Rich et al., 2015; Cazettes et al., 2016). However, the relationship between neuronal coding and perceptual discrimination is unknown because neither the perceptual nor the neuronal discriminability of ITD has been tested as a function of BC.

To investigate this question, we used the pupil dilation response (PDR) to determine how the owl’s ITD discrimination threshold changes with BC. The PDR allows measurement of the behavioral discrimination threshold without extensive training (Bala & Takahashi, 2000). We also determined neuronal ITD discrimination threshold from previous recordings in the midbrain auditory space map and characterized how ITD tuning and variability changed with BC (Cazettes et al., 2016). We show that neuronal ITD tuning resolution is not the sole predictor of perceptual discrimination performance. However, we find that perceptual discrimination is consistent with the average neuronal discrimination threshold in the auditory space map, which is influenced by tuning resolution, firing rate magnitude, and response variability. We now show that modification of a previously proposed habituation model that explained the owl’s discrimination in azimuth and elevation (Bala et al., 2003, 2007) also describes changes in discrimination of ITD as sensory reliability changes. These results extend previous studies and provide a more complete view of the relationship between neural coding and sensory discrimination in the barn owl’s sound localization system.

## Materials and Methods

### Behavior

The subjects were five captive-bred barn owls of either sex. Headplates were surgically attached prior to experimentation, after which the owls were given at least two weeks to recover. Procedures were approved by the Institutional Animal Care and Use Committee of the University of Oregon.

Procedures for measuring the PDR were performed as previously described (Bala & Takahashi, 2000). Stimuli consisted of 100 ms noise bursts with 5 ms rise and fall times, presented at 50 dB SPL. Sounds were presented through earphones (Etymotic ER-2) placed in the ear canal, which were separately attenuated (TDT PA-5). BC was varied by adding independent noise to the sound at one ear. The sound at that ear was a combination of the coherent noise carrying the ITD and the independent noise. The amplitudes of the independent noise and coherent noise were varied so that at BC = 1 the sound was only the coherent noise and at low BC the sound was dominated by the independent noise. The input was scaled to maintain the level of the stimulus at a constant. The BC is the measured cross correlation between the sound signals delivered to the ears. After at least 100 presentations of the habituating stimuli at ITD = 0, test stimuli were presented at one of a few different ITDs. Test stimuli were repeated 4-6 times, separated by 40-50 presentations of the habituating stimuli.

Owls were wrapped in a tight-fitting jacket and laid in the stereotax while headfixed for sessions that lasted no longer than 90 minutes at a time. An infrared pupillometer was placed a few millimeters from one eye, which converted pupil size into voltage. Pupil size was measured for 1 s before and 2 s after sound onset. While the eyelid was held slightly open, the owls could still extend the nictitating membrane to moisten the cornea. Trials were discarded if the pupil was fully or partially obscured by the nictitating membrane during the 2 s gathering period. Custom MATLAB code was used to analyze PDRs. PDR magnitudes were calculated by integrating the voltage over 1 s after sound onset, relative to the mean voltage during 25 ms before and after sound onset. PDR magnitudes were then normalized into z-scores, enabling pooling across sessions varying BC within the same owl. The first 100 habituating trials were excluded from analysis, so that analysis only included fully habituated PDR trials. Data were analyzed using signal detection theory, which enabled us to convert pupil sizes into hit and false alarm rates, which were then used to construct receiver-operating characteristic (ROC) curves. The percent correct could then be calculated by integrating the hit rate over the false alarm rate (integrating the ROC curve) to determine discriminability, with a discrimination threshold set at 75 percent correct.

### Neurophysiology

Neurophysiological data analyzed in this study were collected and partially analyzed in a previous study (Cazettes et al., 2016). Briefly, extracellular recordings of isolated ICx neurons were made from two adult female barn owls under ketamine and xylazine anesthesia. Experiments were performed in a double-walled sound-attenuating chamber. Sounds were delivered over calibrated earphones inserted in the ear canal. ITD tuning was measured by varying ITD in 10 μs steps over 10 repetitions using broadband noise signals (0.5 – 10 kHz) at BC levels ranging from 0 to 1 in 0.1 steps. BC was varied by adding independent noise signals to the sounds at the left and right ears. The value of BC is given by 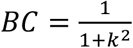, where *k* is the ratio between the root-mean-square amplitudes of the independent noise signals and the time-shifted noise signals that carry the ITD. Detailed experimental methods are described in (Cazettes et al., 2016).

### Neural discriminability

Behavioral ITD discriminability measured using PDR was compared to neural ITD discriminability from responses of single ICx neurons to ITD and BC. The Fisher information *J*(*θ*) places a lower bound to ITD discriminability from neural responses. Specifically, the 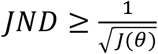 (Seriès et al., 2009). The theory that the sound localization system is designed to maximize information about ITD predicts that the owl’s discrimination threshold approaches the bound placed by the Fisher information. The JND from neural responses was therefore estimated from the Fisher information *J*(*θ*) as 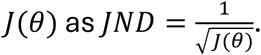.

We computed the linear Fisher information *J*(*θ*) from the neural responses of single Icx neuron as 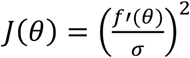 where *f′*(*θ*) is the derivative of the tuning curve and *σ* is the standard deviation of the neural response. The derivative of the tuning curve was approximated by first fitting a Gaussian function to the mean spike count response to estimate the tuning curve *f*(*θ*) and then computing the derivative of the Gaussian function.

### Habituation model

A habituation model was used to examine how responses of midbrain space-specific neurons can lead to the behavioral ITD discrimination results of the PDR experiments. We used a version of the habituation model that previously described results of PDR experiments determining the minimum audible angle in azimuth and elevation (Bala et al., 2003, 2007). The previous model was modified to include the non-uniform distribution of preferred azimuth and elevation that is observed in the midbrain auditory space map (Knudsen, 1982; Fischer and Peña, 2011, 2017) and to include model responses to ITD and BC.

The habituation model proposes that the PDR response is mediated by a population of neurons that habituate to repeated input from midbrain space-specific neurons. The habituating neurons inducing PDR receive topographic input from the midbrain space-specific neurons. The responses of the habituating neurons to a novel stimulus are determined by comparing the responses of space-specific neurons to the novel stimulus to the mean responses of space-specific neurons to the habituating stimulus, relative to the standard deviation of responses of space-specific neurons to the habituating stimulus.

Specifically, let *R*_*SSN,I*_(*n*) denote the response of the *i*^*th*^ space-specific neuron to the stimulus on trial number *n*. For each stimulus type of ITD, azimuth, or elevation, responses of model space-specific neurons were simulated as Poisson random variables with a mean spike count that varied as a Gaussian-shaped function of the stimulus: *R*_*SSN,I*_(*x*) *∼ Poisson*(*λ*(*x*)). The mean rates for the response to azimuth *θ* and elevation ϕ were 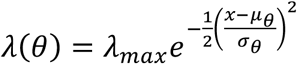 and 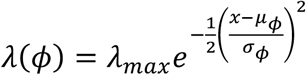, respectively. The preferred azimuth μ_*ϕ*_ was drawn from a Gaussian distribution with mean 0 deg and standard deviation 23.3 deg, and the preferred elevation μ_;_ was drawn from a piecewise-defined function that consists of two Gaussian shaped functions with a common mean of −23 deg and a standard deviation of 12.15 deg for directions less than −23 deg and a standard deviation of 28.6 deg for directions greater than −23 deg, following the non-uniform distribution of preferred azimuth and elevation in the auditory space map (Knudsen, 1982; Fischer and Peña, 2011, 2017). The width parameter of the mean rate function in azimuth was 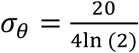 deg and the width parameter in elevation was 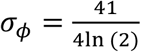 deg to produce tuning curve half widths of 20 degrees in azimuth and 41 degrees in elevation (Bala et al., 2007).

To simulate midbrain space-specific responses to ITD at different BC levels, the stimulus ITD was corrupted with Gaussian noise with a standard deviation that increased exponentially as BC decreased (Fischer and Peña, 2011) and the width of the Gaussian tuning response function increased as BC decreased (Cazettes et al., 2016). The responses of space-specific neurons on each trial were drawn from a Poisson distribution *R*_*SSN,I*_(*ITD, BC*) *∼ Poisson*(*λ*(*ITD, BC*)), where *ITD∼* 𝒩(*ITD*_*STIMULUS*_, *σ*(*BC*)) and the noise standard deviation depended on BC as *σ*(*BC*) = 219.34*e* ^−11.31 *×* BC^ μs (Fischer and Peña, 2011). Note that the noise corrupting the stimulus ITD is shared among all model space-specific neurons and thus introduces noise correlations to the responses. The mean rate for the response to ITD and BC was 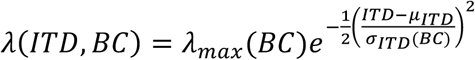. The maximum spike count varied quadratically as a function of BC, *λ*_*max*_(*BC*) = 2.04*BC*^2^ + 2.55*BC* + 0.8, which averages the measured trends in (Albeck and Konishi, 1995) and (Cazettes et al., 2016). The preferred ITD was given by 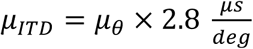 and the width parameter of the Gaussian tuning function in ITD varied exponentially with BC 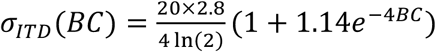 (Cazettes et al., 2016). Here, the multiplicative factor of 2.8 converts degrees azimuth to microseconds ITD, according to the relationship found in barn owl head related transfer functions (Hausmann et al., 2009; Cox and Fischer, 2015). The space-specific neuron responses form the input to the habituating layer of neurons.

Let *R*_*HAB*,0_(*n*) denote the response of the *i*^*th*^ habituating layer neuron to the stimulus on trial *n*. This is given by

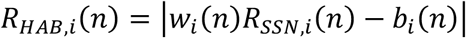

where the weight is the reciprocal of the standard deviation of the midbrain space-specific neuron response 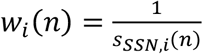 and the bias is the ratio of the mean to the standard deviation of the space-specific neuron response 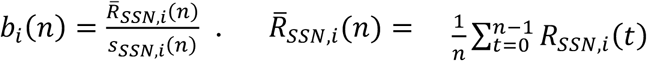 is the average and 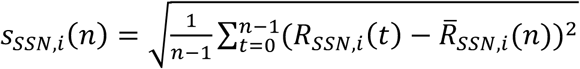 is the standard deviation of the response of the midbrain space-specific neuron over the preceding *n* habituating trials. Therefore, the habituating layer neurons have large responses when the input from space-specific neurons differs from the preceding input, and the deviant response is amplified when the preceding input has been reliable.

It is assumed that discrimination is based on the average activity in the habituating population, 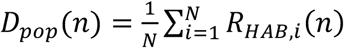. and that the owl detects a change in the stimulus responses when *D*_*pop*_ exceeds a constant threshold. We used the value of *D*_*pop*_ on the test trial to compare the model discriminability to the behavioral discriminability.

## Code Accessibility

Python code to run the simulation is available at https://github.com/brian-fischer/ITD-Discrimination.

## Results

### Behavioral acuity

We first determined the barn owl’s discriminability of ITD at different BC levels. We used the pupil dilation response (PDR) (Bala et al., 2003, 2007) to measure discriminability in five owls. The PDR is a robust measure of discriminability that does not rely on the owl turning its head. The JND for perfectly correlated sounds (BC = 1) ranged from 6 to 14 *μ*s, which is consistent with the previously measured discrimination threshold in azimuth of 3 degrees (Bala et al., 2003). The dependence of the JND on BC was consistent across owls, where the owls showed a small increase in JND as BC decreased to 0.5 and then a large increase for BCs below 0.5 (Figure 1). The dependence of the JND on BC was well described by an exponential curve in each of five owls tested (Figure 1; r > 0.9 for each).

**Figure 1:**
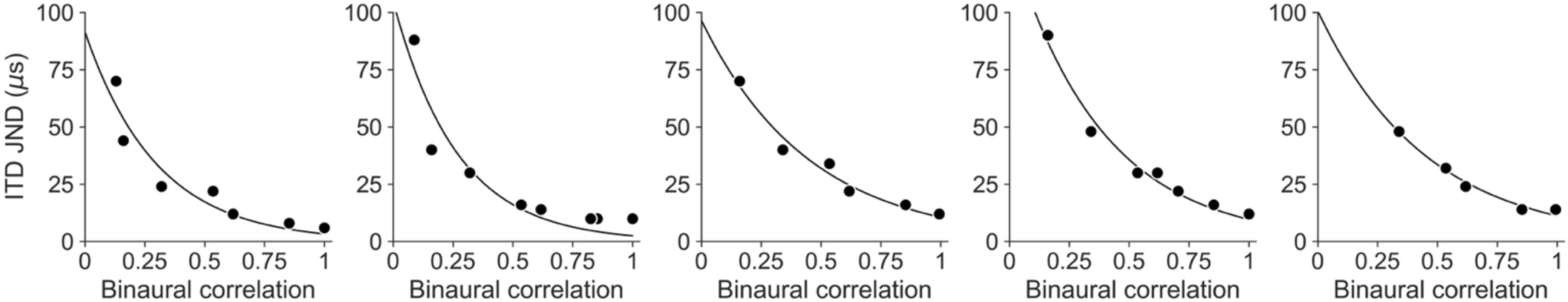
ITD discrimination thresholds. The behavioral just-noticeable-difference (JND) in ITD was computed for a range of binaural correlation (BC) levels in five owls. For each owl the JND vs. BC data were well-fit by an exponential curve (solid line; r > 0.9 for each).

### Neuronal ITD tuning

We then tested whether the behavioral JND is predicted by the spatial resolution of ITD responses of midbrain auditory space map neurons, as was found previously for azimuth and elevation (Bala et al., 2003, 2007). Previous studies demonstrated that the widths of ITD tuning curves of midbrain auditory space map neurons in ICx (Cazettes et al., 2016) and OT (Saberi et al., 1998) increase exponentially as BC decreases. While this exponential pattern is consistent with an exponential relationship between the behavioral ITD JND and BC, is the change in ITD tuning curve width proportional to the change in behavioral ITD JND? In contrast to the prediction of Bala et al. (Bala et al., 2003, 2007), we found that decreasing BC caused a much greater increase in behavioral ITD JND than the increase in the widths of ITD tuning curves in ICx (Figure 2). For example, at BC = 0.6, behavioral ITD JNDs were more than 2 times the JNDs for perfectly correlated sound (BC = 1), whereas ITD tuning widths of ICx neurons only increased by 1.06 times. In fact, average ITD tuning width of ICx neurons never reached twice the tuning width for BC=1. Example cases of single ICx neurons where the ITD tuning width reached double the tuning width found for perfectly correlated sounds only occurred when BC was 0.2 or lower. At these lowest BCs the behavioral ITD JND was between 4 and 12 times the value found for perfectly correlated sounds (7.5 times on average). Thus, the ratio of behavioral ITD JND found at different BC values is not predicted by the ITD tuning widths of ICx neurons at those BCs.

**Figure 2:**
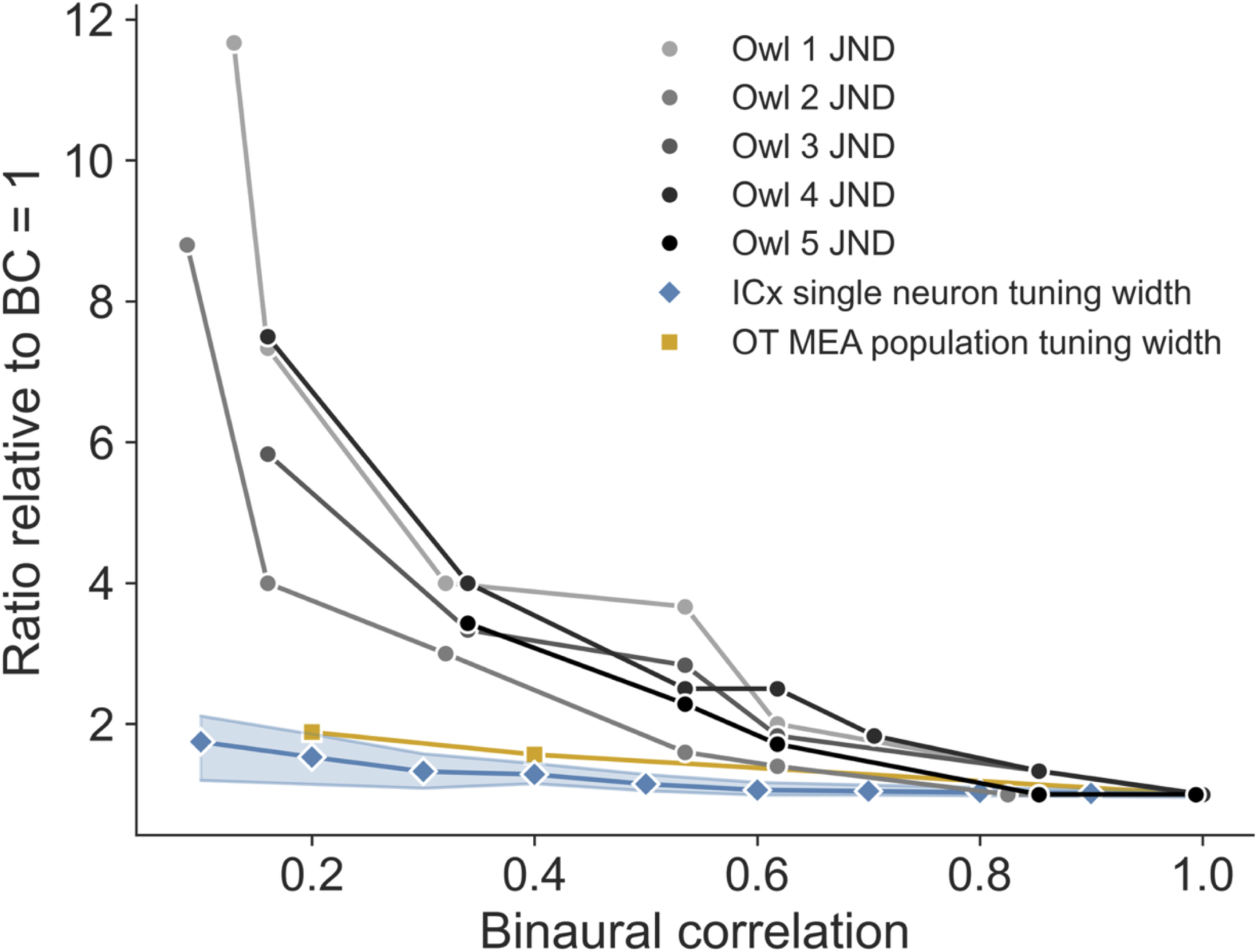
Change in JND and neuronal resolution. The ratio of the behavioral ITD JND at each BC to the JND at BC = 1 for the five owls (greyscale circles). The mean ratio of the single-neuron ITD tuning curve widths for ICx neurons at each BC relative to BC = 1 (blue diamonds). The upper and lower blue lines show the first and third quartiles (adapted from Figure 3D in (Cazettes et al., 2016)). The mean ratio of the OT multi-electrode array (MEA) recorded population activity width at each BC relative to BC = 1 (orange squares) from (Ferger et al., 2021).

In addition to the analysis of single-neuron ITD tuning curves’ width, we examined the tuning width of population activity in the auditory space map in OT at different BC values (Ferger et al., 2021). Ferger et al. (2021) estimated the width of population activity in OT from simultaneously recorded neurons using multielectrode arrays covering wide areas of the OT auditory space map. Consistent with single-neuron responses in ICx, the rate of increase in behavioral ITD JND as BC decreased greatly exceeded the rate of increase in the width of population activity in OT (Figure 2). Thus, contrary to the observation that the ratio of the behavioral JND for elevation and azimuth (with BC = 1) matches the ratio of tuning curve widths for elevation and azimuth in ICx (Bala et al., 2007), the ratio of behavioral ITD JNDs at decreased BCs does not match that of the spatial resolution of ITD tuning in ICx and OT at these same decreased BCs. This difference between behavioral discriminability and neural ITD tuning resolution is expected because decreasing BC causes an increase in sensory noise, and discriminability is dependent on both changes in the mean and the variability of neural responses.

### Neuronal ITD discrimination

We then further compared the behavioral ITD JND to the optimal discriminability of ITD from single ICx neurons (Cazettes et al., 2016). Perhaps, the behavioral ITD JND may correspond to optimal discrimination when both tuning resolution and variability are considered. We used the Fisher information computed from ICx responses to estimate the single-neuron JND (Seriès et al., 2009). The Fisher information was computed as 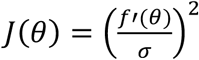 where *f′*(*θ*) is the derivative of the tuning curve and *σ* is the standard deviation of the neural response. The tuning curve was approximated by fitting a Gaussian function to the mean spike count response (Figure 3). The ITD tuning curves were well described by the Gaussian function at each BC, as seen in the examples in Figure 3A,B. The relative error in the fit of the Gaussian function to the mean spike count did not vary significantly with BC (p = 0.11, Kruskal-Wallis, n = 46; Figure 3C).

**Figure 3:**
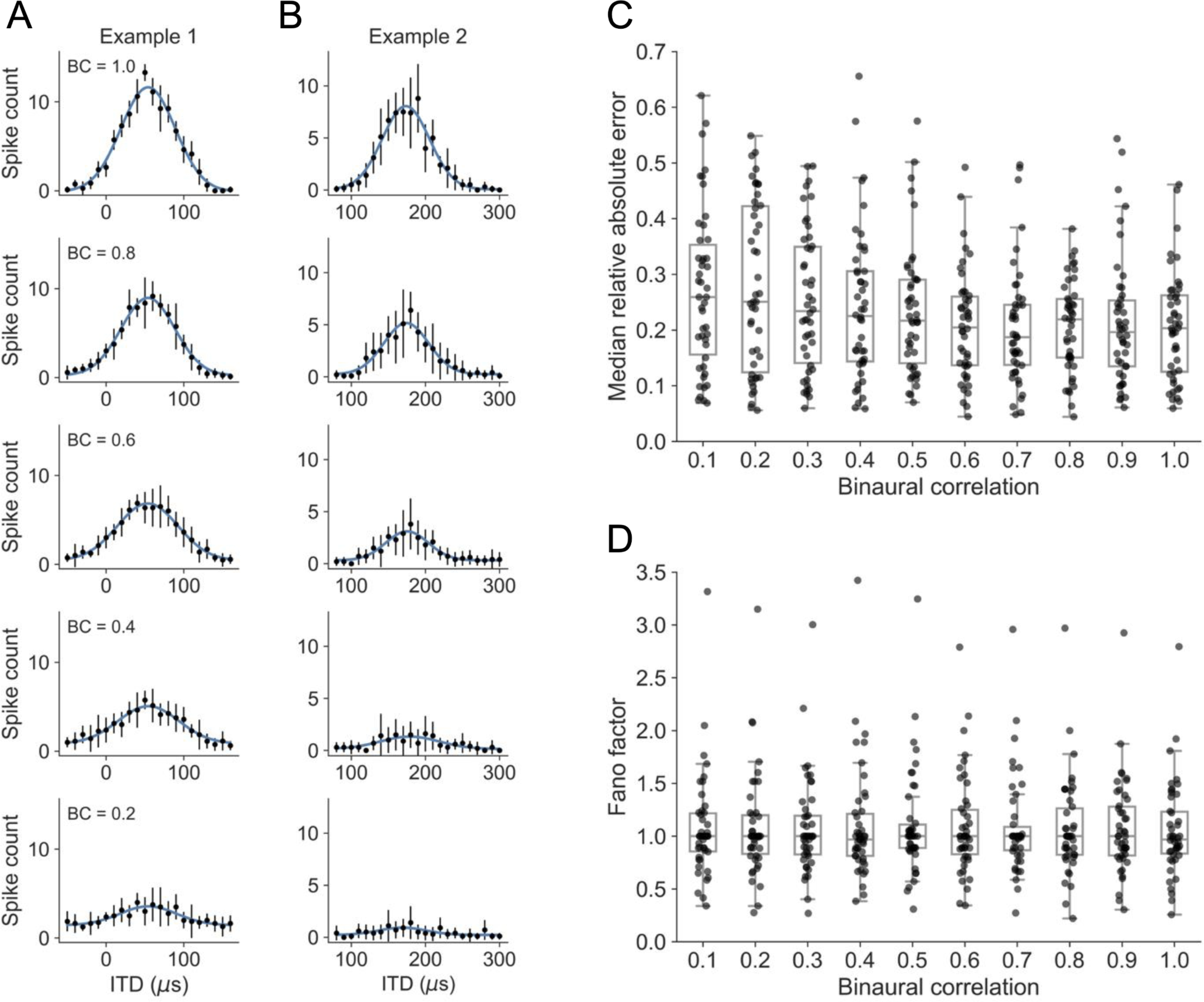
ITD tuning and variability. (A,B) Examples of the mean spike count response (black circles) of ICx neurons to ITD demonstrating the accurate fit with a Gaussian function (blue curve) at different BC values. The error bars represent the standard deviation of the spike count. (C) The median over ITD of the relative absolute error in the fit of the Gaussian function to the mean spike count at each BC. The median relative absolute error did not vary significantly with BC (p = 0.11, Kruskal-Wallis, n = 46). (D) The median over ITD of the Fano factor in ICx responses at each BC. The median Fano factor did not vary significantly with BC (p = 0.99, Kruskal-Wallis, n = 46). The box plots in C and D have boxes that show the first quartile, median, and third quartile, and whiskers that extend to the minimum and maximum values that are not outliers. Outliers are points more than 1.5 times the interquartile range below the first quartile and above the third quartile. The strip plots show individual data points with random jitter in the azimuthal direction to aid visibility.

The responses shown in Figure 3A,B illustrate that the spike count variability changed with the mean spike count. We used the Fano factor, defined as the ratio of the variance to the mean of the spike count, to analyze spike count variability. We observed Fano factors below and above 1, indicating the presence of under- and overdispersion in ICx responses (Figure 3D). The Fano factor did not vary significantly with BC (p = 0.99, Kruskal-Wallis, n = 46), and the overall median value was 1.0.

The Gaussian fits to the ITD tuning curve and the spike count variances were used to compute the Fisher information and then estimate the ITD discrimination performance of ICx neurons. The optimal ITD discriminability of single ICx neurons showed a range of values around the behavioral JND (Figure 4). Similarly to previous results in azimuth and elevation with BC=1 (Bala et al., 2003, 2007), there were ICx neurons with better or worse performance than the behavioral JND over the full range of BC values. Additionally, the median of the JNDs from ICx neuronal responses was consistent with the behavioral JND values. These results are consistent with previous investigations of the relationship between behavioral and neuronal stimulus discriminability showing that behavioral discrimination does not reach the optimal value supported by the most sensitive neurons but is consistent with the average performance of ICx neurons.

**Figure 4:**
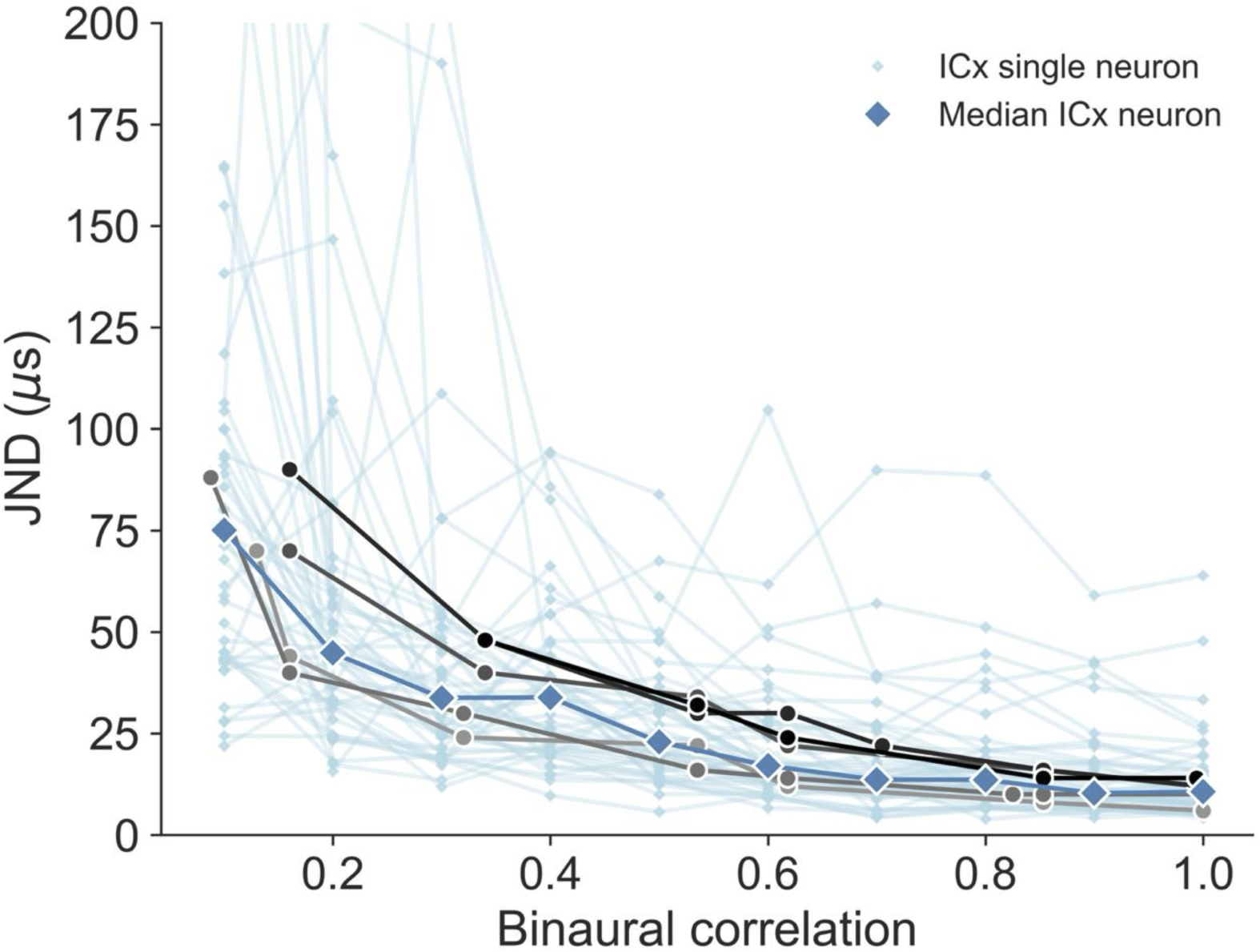
Neuronal discrimination. Comparison of ITD discrimination from behavior (greyscale circles) and single neuron responses in ICx (small light blue diamonds). The large blue diamonds are the median JND over all neurons.

### Habituation model discrimination performance

We then tested whether the behavioral ITD discrimination across BC values can be explained by a habituation model that has been used to explain the behavioral JND for azimuth and elevation at BC=1 (Bala et al., 2003, 2007). The habituation model proposes that the PDR reflects the average activity of a population of neurons that are driven by, and habituate to, activity from the auditory space map. The model assumes that the amount of habituation is higher for activity patterns that are less variable in time. So, habituating layer neurons will have the largest responses to input from the auditory space map that deviate from their recent inputs and when the recent inputs have shown little variability. With this construction, the habituating layer neural activity resembles the magnitude of the z-score of the current auditory space map activity, relative to the mean and standard deviation of previous auditory space map activity. It is assumed that the mean activity of the habituating population determines the JND. Once habituated, the owl will detect a difference in the stimulus when the mean activity of the habituating population crosses a fixed threshold.

We first determined the discrimination threshold so that the model produced responses consistent with behavioral discriminability in azimuth and elevation at BC=1, replicating previous results (Bala et al., 2003, 2007). Here, the model space-specific neurons have tuning curve half widths in elevation (41 deg) that are approximately twice the tuning curve half widths in azimuth (20 deg) (Figure 5A). Correspondingly, the JND in elevation (8.2 deg) is approximately twice the JND in azimuth (3.9 deg) (Figure 5B,C).

**Figure 5:**
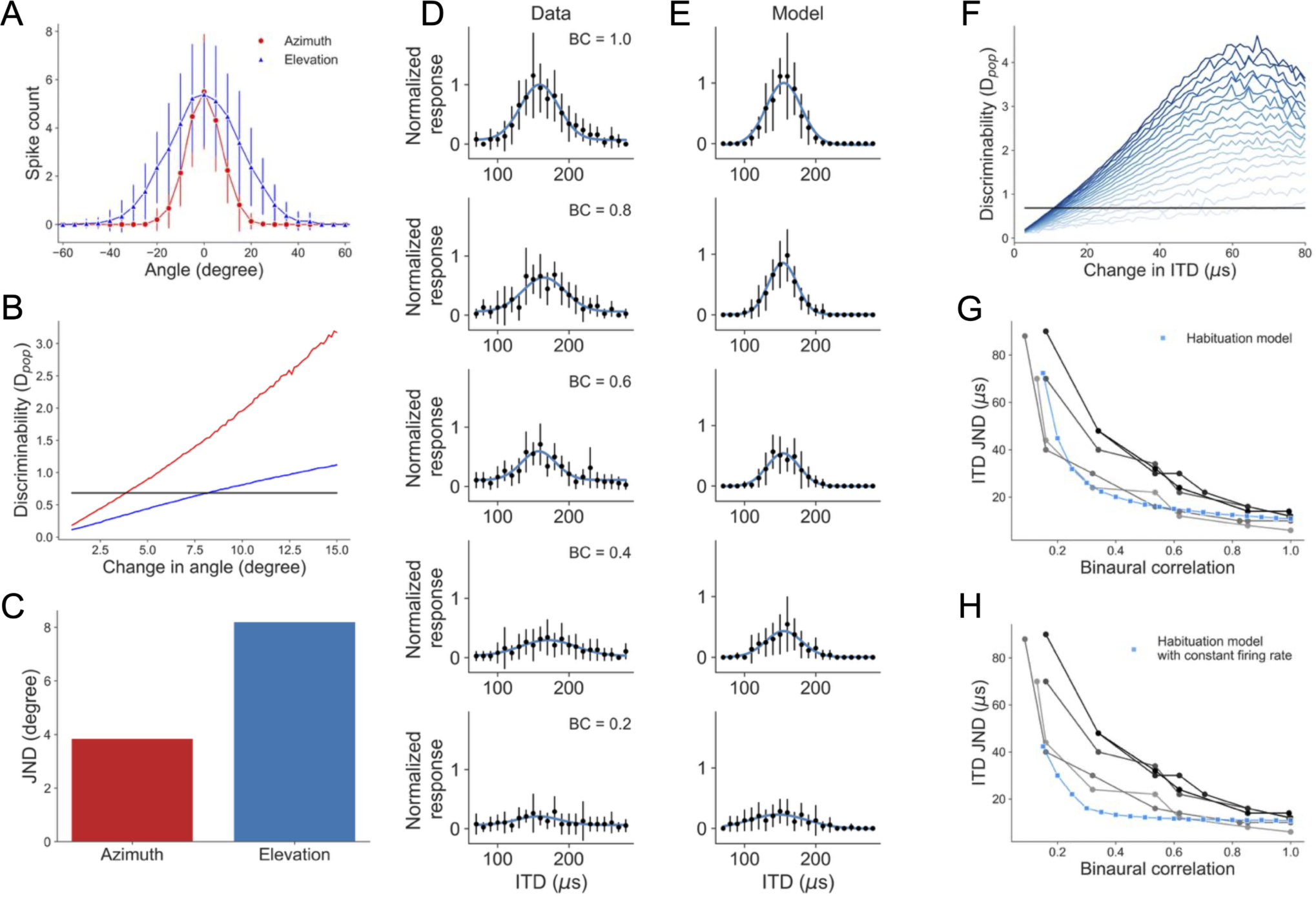
Habituation model. (A) Example tuning curves of model space-map neurons to azimuth (red) and elevation (blue). (B) Discriminability statistic from the habituating population as a function of the angular change in the stimulus for azimuth (red) and elevation (blue). The discrimination threshold (black) was selected to reproduce previously reported results, BC=1 (Bala et al., 2007). (C) Model JND in azimuth and elevation using the discrimination threshold shown in B. (D) Example ITD tuning curves of ICx and (E) a model space-map neuron for different BC values. Mean spike count responses were fit by Gaussian curves (blue). (F) Discriminability statistic from the habituating population as a function of the change in ITD for BC values ranging from low BC (BC = 0.15; light blue) to high BC (BC = 1; dark blue) in steps of 0.05. The discrimination threshold is shown in the black line. (G) Comparison of ITD discrimination from behavior (greyscale circles) and the habituating neuron model (blue squares). (H) Comparison of ITD discrimination from behavior (greyscale circles) and the habituating neuron model when the firing rate of model neurons is constant at the value for BC = 1 (blue squares).

We modified the habituation model of (Bala et al., 2003, 2007) to allow for model space-specific neurons to respond to ITD *and* BC. Model space-specific neurons were driven by a stimulus ITD that was corrupted by random Gaussian noise with a standard deviation that increased exponentially as BC decreased (Fischer and Peña, 2011). The tuning widths increased and firing rates decreased as BC decreased, consistent with experimental observations (Figure 5D,E) (Albeck and Konishi, 1995; Saberi et al., 1998; Cazettes et al., 2016; Ferger et al., 2021). Model space-specific neuron spike counts had a Poisson distribution, consistent with the observation that the median Fano factor for ICx neurons was one (Figure 3D).

We tested the model’s discrimination of ITD at different BC levels using the discrimination threshold found (at BC=1) so that the model produced responses consistent with behavioral discriminability in azimuth and elevation. This model was consistent with the owls’ dependence of JND on BC (Figure 5F,G). The consistency of the model with behavior under decreasing BC depends on a decrease in space map firing rates, and thus a decrease in spike count variability. When firing rates of model space map neurons were held constant at their maximum value (BC=1), the model performance exceeded the owl’s discrimination performance, decreasing more slowly with BC (Figure 5H). The model shows that a habituation model can explain the owl’s discrimination in azimuth, elevation, and ITD at different BCs. Moreover, these results show that both response resolution and variability of midbrain auditory space map neuronal responses can determine the owl’s discrimination of sound location.

## Discussion

ITD reliability describes how corruptible the ITD cue for sound source localization is when faced with concurrent sounds (Cazettes et al., 2014). In the barn owl, ITD reliability derives from the responses of interaural phase difference-tuned neurons in the inferior colliculus and their target ITD-tuned neurons in the ICx (Cazettes et al., 2014). We studied the relationship between ITD-sensitive neuronal responses and behavioral ITD-discrimination while presenting varying levels of binaural correlation, which directly changed ITD reliability (Licklider, 1948; Jeffress et al., 1962). We show that perceptual sensitivity to ITD degrades exponentially as binaural correlation decreases. This decline in behavioral performance, however, could not be accounted for by concomitant changes in neuronal tuning width alone. ITD tuning broadened in ICx neurons with decreasing BC, however, the rate of broadening was insufficient to explain the exponential increase in behavioral JNDs. Behavioral JNDs were more closely aligned with the average ITD discriminability predicted by Fisher information from single-unit ICx responses, thus integrating tuning resolution, response magnitude, and response variability. This suggests that perceptual acuity reflects the average information content across the neural population, rather than the optimal performance of the most sensitive neurons.

A habituation model based on that of (Bala et al., 2003, 2007) provided additional support for this interpretation. Only after incorporating empirically observed changes in tuning width, firing rate, and response variability did the model accurately capture behavioral ITD discrimination across BC levels. In contrast, when model firing rates were held constant, performance deviated from behavioral data, underscoring the importance of reduced response magnitude and increased variability limiting sensory discriminability in degraded sensory environments. Additionally, decreasing binaural correlation resulted in linked decreases in both mean neuronal spike count and spike count variability, with a median Fano Factor near one, suggesting that regulation of spike count variability is important for behavioral discrimination.

This work advances our understanding of the relationship between neural coding and perceptual stimulus discrimination. We provide additional support to the previous conclusion that the response properties of midbrain auditory space maps in ICx and OT determine the ability to discriminate changes in sound location by incorporating the influence of changes in sensory reliability in our understanding of how midbrain auditory space maps in ICx and OT determine the ability to discriminate changes in sound location. We showed that a habituating neuron model can explain how ICx responses to azimuth, elevation, and ITD as BC changes are translated into perceptual stimulus discrimination limits. This provides a more comprehensive understanding of how neural coding in the midbrain auditory space map is related to sound location perception.

## References

Abbott LF, Dayan P (1999) The effect of correlated variability on the accuracy of a population code. Neural Comput 11:91–101.

Albeck Y, Konishi M (1995) Responses of neurons in the auditory pathway of the barn owl to partially correlated binaural signals. J Neurophysiol 74:1689–1700.

Averbeck BB, Latham PE, Pouget A (2006) Neural correlations, population coding and computation. Nat Rev Neurosci 7:358–366.

Bala AD, Takahashi TT (2000) Pupillary dilation response as an indicator of auditory discrimination in the barn owl. J Comp Physiol [A] 186:425–434.

Bala ADS, Spitzer MW, Takahashi TT (2003) Prediction of auditory spatial acuity from neural images on the owl’s auditory space map. Nature 424:771–774.

Bala ADS, Spitzer MW, Takahashi TT (2007) Auditory spatial acuity approximates the resolving power of space-specific neurons. PLoS ONE 2:e675.

Cazettes F, Fischer BJ, Pena JL (2014) Spatial cue reliability drives frequency tuning in the barn Owl’s midbrain. eLife 3:e04854.

Cazettes F, Fischer BJ, Peña JL (2016) Cue reliability represented in the shape of tuning curves in the owl’s sound localization system. J Neurosci 36:2101–2110.

Cox W, Fischer BJ (2015) Optimal prediction of moving sound source direction in the owl. PLoS Comput Biol 11:e1004360.

Ferger R, Shadron K, Fischer BJ, Peña JL (2021) Barn owl’s auditory space map activity matching conditions for a population vector readout to drive adaptive sound-localizing behavior. J Neurosci 41:10305–10315.

Fischer BJ, Peña JL (2011) Owl’s behavior and neural representation predicted by Bayesian inference. Nat Neurosci 14:1061–1066.

Fischer BJ, Peña JL (2017) Optimal nonlinear cue integration for sound localization. J Comput Neurosci 42:37–52.

Harper NS, McAlpine D (2004) Optimal neural population coding of an auditory spatial cue. Nature 430:682–686.

Hausmann L, von Campenhausen M, Endler F, Singheiser M, Wagner H (2009) Improvements of sound localization abilities by the facial ruff of the barn owl (Tyto alba) as demonstrated by virtual ruff removal. PloS One 4:e7721.

Jeffress LA, Blodgett HC, Deatherage BH (1962) Effect of interaural correlation on the precision of centering a noise. J Acoust Soc Am 34:1122–1123.

Kettler L, Griebel H, Ferger R, Wagner H (2017) Combination of interaural level and time difference in azimuthal sound localization in owls. eNeuro 4.

Knudsen EI (1982) Auditory and visual maps of space in the optic tectum of the owl. J Neurosci Off J Soc Neurosci 2:1177–1194.

Licklider JCR (1948) The influence of interaural phase relations upon the masking of speech by white noise. J Acoust Soc Am 20:150–159.

Moiseff A (1989) Bi-coordinate sound localization by the barn owl. J Comp Physiol [A] 164:637–644.

Moreno-Bote R, Beck J, Kanitscheider I, Pitkow X, Latham P, Pouget A (2014) Information-limiting correlations. Nat Neurosci 17:1410–1417.

Peña JL, Konishi M (2004) Robustness of multiplicative processes in auditory spatial tuning. J Neurosci 24:8907–8910.

Poganiatz I, Nelken I, Wagner H (2001) Sound-localization experiments with barn owls in virtual space: influence of interaural time difference on head-turning behavior. J Assoc Res Otolaryngol JARO 2:1–21.

Pouget A, Dayan P, Zemel RS (2003) Inference and computation with population codes. Annu Rev Neurosci 26:381–410.

Rakerd B, Hartmann WM (2010) Localization of sound in rooms. V. Binaural coherence and human sensitivity to interaural time differences in noise. J Acoust Soc Am 128:3052–3063.

Rich D, Cazettes F, Wang Y, Peña JL, Fischer BJ (2015) Neural representation of probabilities for Bayesian inference. J Comput Neurosci 38:315–323.

Saberi K, Takahashi Y, Konishi M, Albeck Y, Arthur BJ, Farahbod H (1998) Effects of Interaural Decorrelation on Neural and Behavioral Detection of Spatial Cues. Neuron 21:789–798.

Seriès P, Stocker AA, Simoncelli EP (2009) Is the Homunculus “aware” of sensory adaptation? Neural Comput 21:3271–3304.

Seung HS, Sompolinsky H (1993) Simple models for reading neuronal population codes. Proc Natl Acad Sci U S A 90:10749–10753.

Trahiotis C, Bernstein LR, Akeroyd MA (2001) Manipulating the “straightness” and “curvature” of patterns of interaural cross correlation affects listeners’ sensitivity to changes in interaural delay. J Acoust Soc Am 109:321–330.

